# The Progenetix oncogenomic resource in 2021

**DOI:** 10.1101/2021.02.15.428237

**Authors:** Qingyao Huang, Paula Carrio-Cordo, Bo Gao, Rahel Paloots, Michael Baudis

## Abstract

In cancer, copy number aberrations (CNA) represent a type of nearly ubiquitous and frequently extensive structural genome variations. To disentangle the molecular mechanisms underlying tumorigenesis as well as identify and characterize molecular subtypes, the comparative and meta-analysis of large genomic variant collections can be of immense importance. Over the last decades, cancer genomic profiling projects have resulted in a large amount of somatic genome variation profiles, however segregated in a multitude of individual studies and datasets. The Progenetix project, initiated in 2001, curates individual cancer CNA profiles and associated metadata from published oncogenomic studies and data repositories with the aim to empower integrative analyses spanning all different cancer biologies.

During the last few years, the fields of genomics and cancer research have seen significant advancement in terms of molecular genetics technology, disease concepts, data standard harmonization as well as data availability, in an increasingly structured and systematic manner. For the Progenetix resource, continuous data integration, curation and maintenance have resulted in the most comprehensive representation of cancer genome CNA profiling data with 138’663 (including 115’357 tumor) CNV profiles. In this article, we report a 4.5-fold increase in sample number since 2013, improvements in data quality, ontology representation with a CNV landscape summary over 51 distinctive NCIt cancer terms as well as updates in database schemas, and data access including new web front-end and programmatic data access. Database URL: progenetix.org

## Introduction

Copy number aberrations (CNA) are present in the majority of cancer types and exert functional impact in cancer development [1, 2]. As understanding cancer biologies remains one of the main challenges in contemporary medical and life sciences, the number of studies addressing genomic alterations in malignant diseases continues to grow. Progenetix is a publicly accessible cancer genome data resource (*progenetix.org*) that aims to provide a comprehensive representation of genomic variation profiles in cancer, through providing sample-specific CNA profiles and associated metadata as well as services related to data annotation, meta-analysis and visualization. Originally established in 2001 with a focus on data from chromosomal Comparative Genomic Hybridization (CGH) studies [3], the resource has progressively incorporated data from hundreds of publications reporting on molecular-cytogenetic (CGH, genomic arrays) and sequencing (whole genome or whole exome sequencing; WGS, WES) based genome profiling experiments. Since the last publication dedicated to the Progenetix resource in 2014 [4], changes in content and features of the data repository and its online environment have vastly expanded its scope and utility to the cancer genomics community. For data content, additions include the complete incorporation of the previously separate arrayMap data collection [5, 6] and of datasets from external resources and projects such as The Cancer Genome Atlas (TCGA; [7, 8]) or cBioPortal [9], as well as the recurrent collection and re-processing of array-based data from NCBI’s GEO or EMBL-EBI’s ArrayExpress [10, 11]. Additionally, data content updates have followed the previous methodology of publication-based data extraction where feasible. Beyond the data expansion, a tight integration with projects of the Global Alliance for Genomics and Health (GA4GH [12]) and ELIXIR - such as serving for implementation-driven development of the Beacon API [13] - has led to an extension of the resource’s features as well as adoption and promotion of emerging open data standards.

Here we present the latest updates on data content, structuring, standardization, access and other modifications made to the Progenetix resource.

## Data expansion and new features

### Genomic profiling data

Over the last two decades, thousands of cancer genomes studies have used the Gene Expression Omnibus (GEO; [14]) for deposition of data from array-based experiments. Data from GEO contributes a substantial fraction of the genomic screening data in the Progenetix collection and has again been expanded in both number of samples and represented platforms. Additionally, we systematically included suitable data from three more resources: ArrayExpress[15], cBioPortal (cBP)[16] and The Cancer Genome Atlas (TCGA)[17] project. As in the previous database updates, we have also included data directly derived from publication supplements and from collaborative projects. Table 1 shows statistics of samples within each resource. Table 2 reports the overall data growth and sample counts stratified by cancer loci since the last update. [4].

**Table 1.** Statistics of samples from various data resources

| Data Source | GEO | ArrayExpress | cBioPortal | TCGA | Total |
| --- | --- | --- | --- | --- | --- |
| No. Studies | 898 | 51 | 38 | 33 | 1’939 |
| No. samples | 63’568 | 4’351 | 19’712 | 22’142 | 138’663 |
| Tumor | 52’090 | 3’887 | 19’712 | 11’090 | 115’357 |
| Normal | 11’478 | 464 | 0 | 11’052 | 23’306 |
| Classifications |  |  |  |  |  |
| ICD-O (Topography) | 100 | 54 | 88 | 157 | 209 |
| ICD-O (Morphology) | 246 | 908 | 265 | 140 | 491 |
| NCIt | 346 | 148 | 422 | 182 | 788 |
| Collections |  |  |  |  |  |
| Individuals | 63’568 | 4’351 | 19’712 | 10’995 | 127’549 |
| Biosamples | 63’568 | 4’351 | 19’712 | 22’142 | 138’663 |
| Callsets | 63’568 | 4’351 | 19’712 | 22’376 | 138’930 |
| Variants | 5’514’126 | 118’4170 | 1’778’096 | 2’654’065 | 10’716’093 |

**Table 2.** Data growth by cancer loci

| Cancer loci | No. in 2014 | No. in 2021 |
| --- | --- | --- |
| Hematopoietic and reticuloendothelial systems | 5269 | 18482 |
| Lymph nodes | 2345 | 5988 |
| Breast | 2271 | 15790 |
| Cerebellum | 1439 | 3465 |
| Brain, NOS | 1342 | 6608 |
| Cerebrum | 1201 | 1712 |
| Liver | 1180 | 3237 |
| Stomach | 1155 | 3176 |
| Skin | 1073 | 3343 |
| Connective, subcutaneous and other soft tissues | 1058 | 2526 |
| Kidney | 1018 | 3617 |
| Colon | 1001 | 5182 |
| Ovary | 733 | 3963 |
| Prostate gland | 735 | 4485 |
| Lung and bronchus | 699 | 10321 |
| Nervous system, NOS | 667 | 926 |
| Urinary bladder | 587 | 1961 |
| Cervix uteri | 529 | 1331 |
| Peripheral nerves incl. autonomous | 523 | 1479 |
| Esophagus | 454 | 1890 |
| Pancreas | 426 | 1620 |
| Thyroid gland | 404 | 1260 |
| Heart, mediastinum, and pleura | 383 | 771 |
| Bones, joints and articular cartilage | 350 | 1205 |
| Spleen | 278 | 636 |
| Other | 4522 | 16268 |
| Total | 31642 | 115359 |

The “ArrayExpress Archive of Functional Genomics Data”, hosted by European Molecular Biology Laboratory - European Bioinformatics Institute (EMBL-EBI), stores functional genomics data submitted by research groups and projects. In this update, we have incorporated the cancer-related genomic profiles which do not have corresponding GEO entries using our analysis pipeline. Overall, data from ArrayExpress added 3’887 samples from 44 projects which resolve to 143 distinct cancer types according to NCIt. Similar to the GEO data acquisition procedure, we have used a combination of text mining methods and expert curation for annotation of technical metadata and biomedical parameter.

The “cBioPortal for Cancer Genomics” is an open-access resource for cancer genomics data, representing different types of molecular screening data from 19’712 samples, derived from 38 studies and mappable to 422 NCIt cancer types. The largest part of genomic data is based on whole exome sequencing analyses from the MSK-TARGET [18] pipeline, with CNA data accessed directly as segment files in genome version hg19/GRCh37. The data was converted into GRCh38 with the *segment-liftover* tool [19], and oncology classifications as well as relevant clinical data were incorporated into our database.

The Cancer Genome Atlas project provides a set of multi-omics data with extensive structured meta-data annotation for a large collection of cancer types, currently through NCBI’s Genomic Data Commons Data Portal (https://portal.gdc.cancer.gov). In this update, we incorporated its CNV profiling data as well as transformed the relevant clinical information into our system (Figure 1).

**Fig 1.**
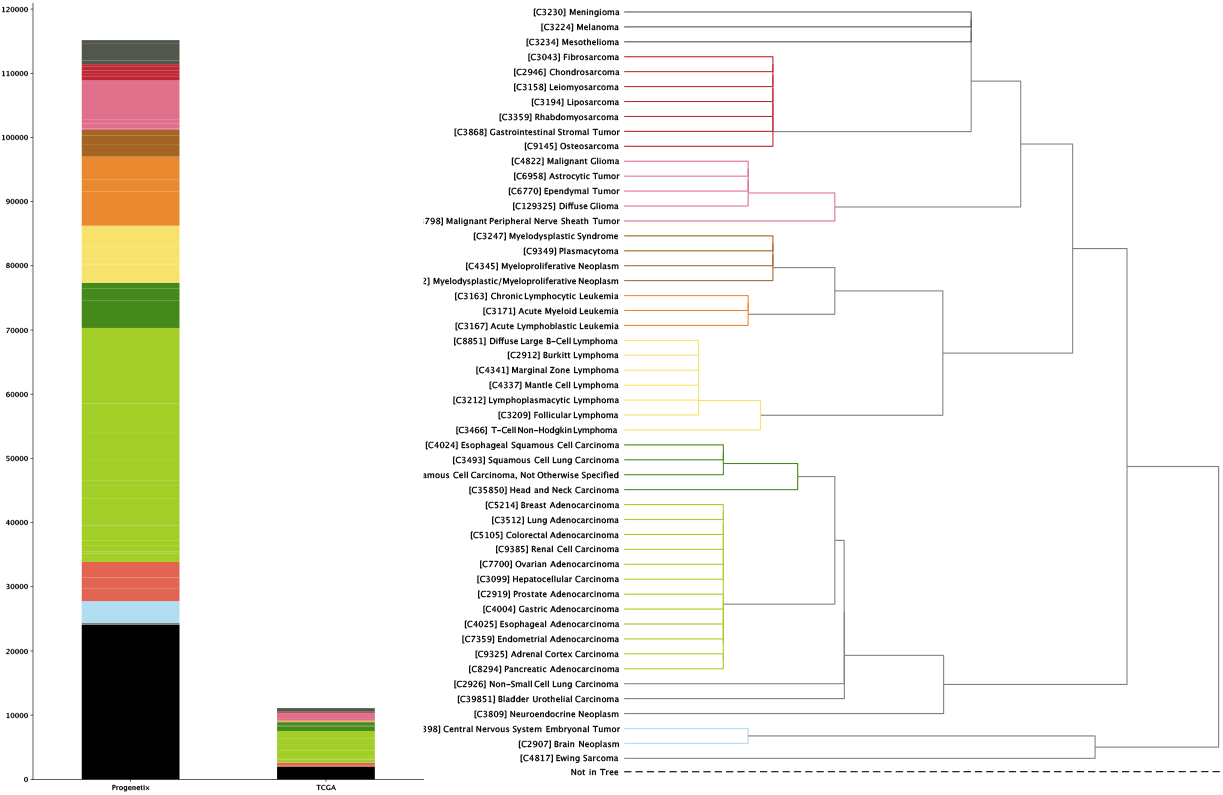
The currently available CNA data points in Progenetix and TCGA. Progenetix database contains 115’357 cancer samples with 92’307 mapped to the 51 defined critical nodes in NCIt ontology tree and 23’050 samples not mapped to the tree (black); whereas TCGA repository contains 11’090 samples with 9’103 samples mapped and 1987 sample not mapped to the tree (black). Colors of the stacked barplot (left) match the branch colors on NCIt ontology tree (right)

### Data processing update

Genomic profiling data in Progenetix originates from a large number of studies which are based on different molecular-cytogenetic and sequencing based technologies. In order to maximize qualitative homogeneity of the final CNA calls, we prefer to download source files with the least amount of pre-processing and apply our in-house data processing pipeline from the arrayMap project [5]. Currently, our analysis workflow handles the raw data based processing for 13 Affymetrix SNP array platforms, including 9 genome-wide arrays - 10K (GPL2641),50K (Hind240 and Xba240) (GPL2004 and GPL2005), 250K (Nsp and Sty) (GPL3718 and GPL3720), Genome-wide SNP (5.0 and 6.0) (GPL6894 and GPL6801, respectively), CytoScan (750K and HD) (GPL18637 and GPL16131) arrays (GPL-prefixed platform coding in brackets according to GEO standard) - as well as the 4 cancer-specific “Oncoscan” arrays - GPL18602, GPL13270, GPL15793 and GPL21558.

### Allele-specific copy number variation

For the subset of SNP array based experiments - where the status of both alleles can be evaluated separately - we have analyzed allele-specific copy number data (ASCN) and incorparated 35’897 LOH profiles into the database. ASCN potentiates new analysis on the same samples. First, probe-wise it gives an overview of germline variant landscape, as used in determining the ancestry background. Second, it allows detection of loss of heterozygosity (LOH) events, including of copy-number neutral event (CN-LOH) which e.g. can be commonly observed in haematological malignancies due to a selective process for duplication of minor disease-prone germline alleles [20, 21]. Lastly, it acts as a second reference for CNA to combat the variability caused by known wave artefacts from array technologies [22]. For all SNP arrays, we have implemented a pipeline to determine probe-wise B-allele frequency (BAF) of SNP probes and perform subsequent segmentation [23, 24]. Subsequently, we use ASCN to assess ancestry provenance of the samples[25] and store the LOH regions of the samples in our genomic variants database.

## Meta-data updates

Over the last years, a number of initiatives have addressed the lack of - or inconsistencies in - data standards for the representation and exchange of technical and “bio”-metadata in life sciences. Since 2016, the Progenetix resource has served as test bed for the implementation-driven design of metadata schemas and data exchange protocols, in conjunction with GA4GH workstreams and driver projects. For the structure and content of the Progenetix resource, this involvement led to the forward-looking adoption of hierarchical ontology classes as a core concept for a robust annotation of categorical metadata such as ‘biocharacteristics’ (e.g. phenotypes, disease categories) as well as the use of registered identifiers for external references using a CURIE syntax. Importantly, the resource informed the development and implemented the principles of a robust hierarchical data model for genomic data collections and exchange formats.

### NCIt ontology mapping

Since its establishment, Progenetix has made use of the *International Classification of Diseases in Oncology,* 3rd Edition (ICD-O 3) [26] for cancer sample classification. While the combination of the ICD-O Morphology and Topography coding systems depicts diagnostic entities with high specificity, the current ICD-O is limited in its representation of hierarchical concepts and does not easily translate to modern ontologies. In comparison, the National Cancer Institute Thesaurus (NCIt; access through http://bioportal.bioontology.org/ontologies/NCIT) is a dynamically developed hierarchical ontology which empowers layered data aggregation and transfer between classification systems and resources. However, due to the comparatively recent development and ongoing expansions, NCIt terms are rarely used in primary sample annotations. In the recent Progenetix update, we performed a data-driven generation of ICD-O - NCIt mappings and added the derived NCIt codes to all (existing and new) samples, to take advantage of NCIt’s hierarchical structure for data retrieval, analysis and exchange (Figure 4B).

### Data summary based on the NCIt hierarchy tree

All cancer samples in Progenetix have been annotated with an NCIt code, resulting in currently 789 distinct NCIt terms. However, as the definition of increasingly specific NCIt terms outruns their incorporation into the hierarchical tree, 98 of these terms so far are not represented in the tree hierarchy. For better illustration, we define 51 prominent nodes under which we summarize and visualize the data collection, where the disease types are both conceptually distinctive (e.g. “carcinoma” as category is too broad and thus its child nodes will be used) and include a considerable number of samples under the term or its child terms. This brings about additional 324 (60 in TCGA) terms not mappable to the selected nodes, resulting in 23,050 (1987 for TCGA) samples excluded from the summary tree counts (black bar in left panel of Figure 1). For terms with multiple occurrences in the tree we define the preferred path to the selected node by prioritizing morphology-based separation. The sample collection in Progenetix as compared to TCGA is summarized with reference to the NCIt coding system (Figure 1; Supplementary Table 1).

### CNV data content by cancer type

With cancer genomes grouped in the 51 NCIt nodes, we assessed their differences in the CNV landscape. The fraction of genome with a copy number alteration (CNV fraction) varies widely among the cancer types (Figure 2; Supplementary Figure 1). Among the most studied cancer types, the breast carcinoma shows a consistent CNV profile as an earlier analysis with frequent chr1q,8q,16p,17q,20 gain and 8p,16q,17p,18,22q loss [27]; the CNV pattern in cervical (chr3 gain), colorectal(chr7, 8q, 13, and 20q gain and 8p, 17p, and 18 loss) carcinoma also correspond with previous observation[28]; similarly for T-cell non-Hodgkin lymphoma [29] and myelodysplastic syndrome [30]. In addition, the genome-wide LOH profile also shows distinction among the cancer types in evaluation (42 out of 51 with at least 20 samples; Supplementary Figure 2). LOH profile of a cancer genome complements its CNV profile with the information of allelic loss. We highlight here a few prominent patterns which have been previously reported: chr3p and 9 in esophageal squamous cell carcinoma [31, 32]; chr18q in colorectal carcinoma [33]; chr13q, 16q and 17p in hepatocellular carcinoma [34].

**Fig 2.**
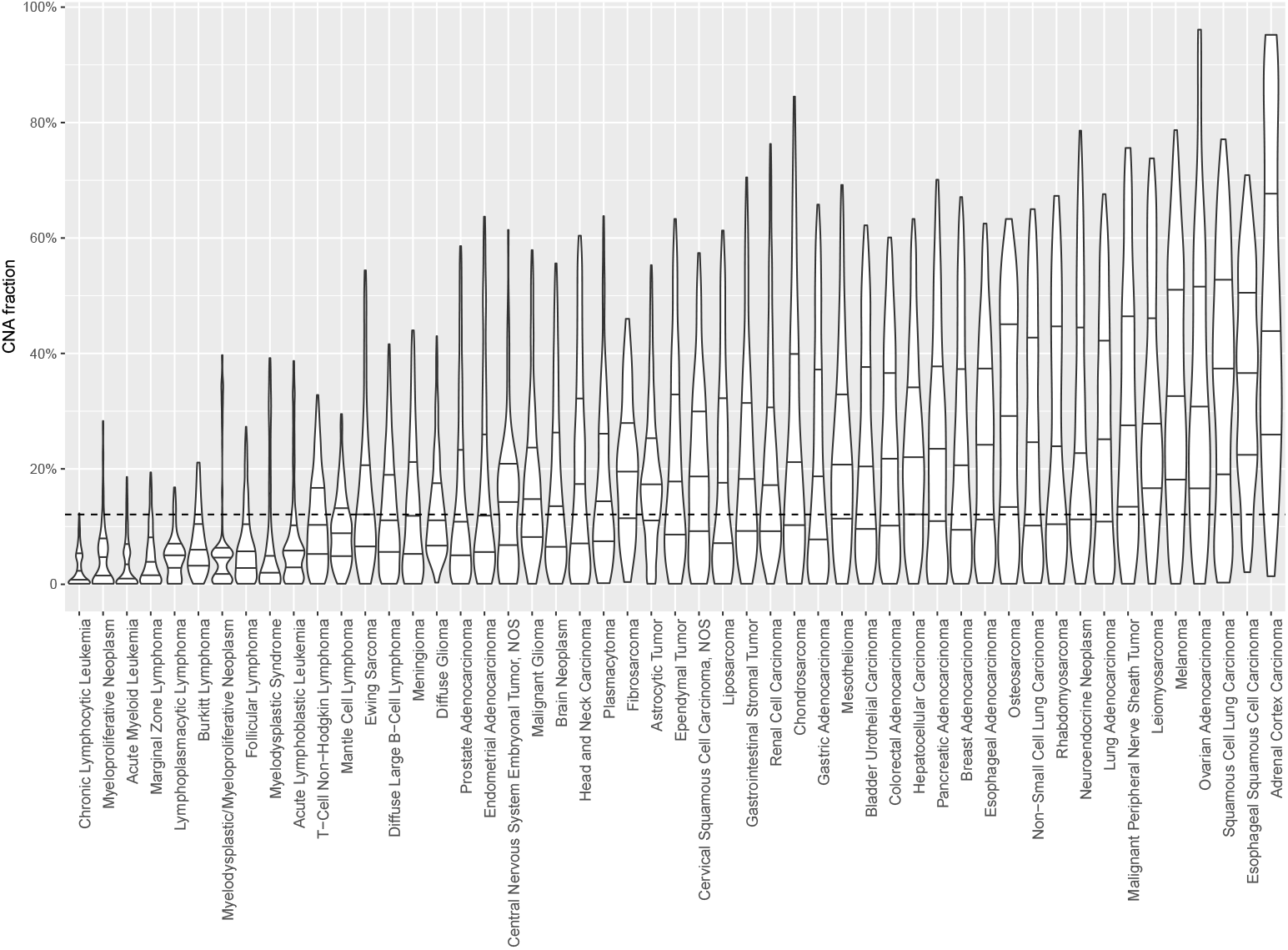
Distribution of genomic CNV fraction in 51 NCIt umbrella nodes. illustrated with violin plot, showing median, first and third quantile. Dashed line indicates the global median CNV fraction at 12.1%. Each category contains between 104 and 11804 CNV profiles (median 904; See Supplementary Table 1).

### Uberon anatomy ontology

While the ICD-O topography system provides organ and substructure specific mapping rooted in traditional clinical and diagnostic aspects of a “tumor entity”, *UBERON* is a cross-species anatomical structural ontology system closely aligned with developmental processes[35]. Its relationship structure allows integrative queries linking multiple databases (e.g. Gene Ontology[36], Protein Ontology[37]), description logic query within the same organism (linking related organs) and between model animals and humans. In this resource update, we have mapped all existing ICD-O T codes to *UBERON* terms, and additionally have provided those as part of the *Monarch* initiative[38], with our latest mapping table made available through a Github repository *progenetix/icdot2uberon*).

### Provenance by geography

As part of the curated metadata provided in the sample representation, we have included geographic point coordinates for each individual sample. As this information is often missing from individual sample annotations, we previously have applied a mapping procedure to assign the samples’ approximate geographic origins. For samples with the submitter’s contact available from repository entries a default point-location in the corresponding city was used; otherwise that of the corresponding author of the associated publication was used. Associated publications were also explored for more detailed descriptions of sample origin. Point coordinates for each city were obtained using the external geographic database GeoNames (www.geonames.org), as detailed previously [39].

### Provenance by ancestry group

While providing a good approximation for the geographic origin of cancer profiling data, which can e.g. be useful for epistemic validation and decision processes, the geographic location of the studies provides limited specificity regarding individual sample provenance, especially when assessing correlations between genomic variants and ancestral population background. Beyond the scope of high-penetrance variants like mutations in the BRCA1/2[40, 41], or RB genes [42] in cancer predisposition, other studies have asserted an influence of genetic background on tumor development[43, 44, 45, 46]. Previously we have developed a method for deriving ancestry groups from un-masked germline variants in cancer genomes, based on reference populations studied in the 1000 Genomes Project [25]. For samples in Progenetix with accessible SNP data, population groups were assigned based on the reference categories mapped to Human Ancestry Ontology (HANCESTRO) terms (Supplementary Table 2). Where available, the respective data is now represented under the “populations provenance” schema for the corresponding biosample entries.

## Updated data access modalities

Since the last release, we have adopted the GA4GH data schema standards and migrated to Phenopackets[47]-formatted response delivery with modified data access points in the user interface. Information about application programming interface (API) methods are provided through the documentation pages (https://info.progenetix.org/categories/API).

### Data standards

In many genomic repositories, databases are structured around experimental outcomes (e.g. variants from a DNA sequencing experiments as collections of VCF files). Recent attempts in evaluating sensible meta-schemas for the representation of genomic variants and related biological or technical metadata, especially with respect to empowering data federation over flexible, networked resources, have led to a set of emerging meta-models and data schemas[48]. The data storage and representation models for the Progenetix resource have been designed to comply with concepts developed by the previous GA4GH Data Working Group [12, 49] and subsequent GA4GH work streams, documented e.g. by the *SchemaBlocks* {S} initiative (http://schemablocks.org). One of the core concepts is the “individual - biosample(s) - variants” meta-model which is applicable to cancer-related analyses with potentially multiple samples representing different stages in the course of disease as well as the underlying genomic background. This hierarchical model provides a solid representation and connection between the physical source of the data and the logical genotyping information and adapts to various scenarios for data aggregation and analysis.

### User interface

The completely re-designed user interface provides flexibility and versatility in query parameters and types and optimized the response delivery. Technically, the query interface for retrieval of sample specific data is built on top of a forward-looking implementation of the GA4GH Beacon API [13] with features from the upcoming version 2 of this standard.

Figure 3 shows the current web interface to perform a CNA query with start and end position range with filter options for cancer type, tissue location, morphology, cell line or geographic location. The top panel of the result page shows a summary with the number of matched samples, variants, calls and the frequency of alleles containing the CNA (Figure 3E). The “Phenopackets” link returns a json document of biosamples with the phenopacket-formated response. The “UCSC region” links externally to a UCSC browser track providing an overview of the genomic elements which map to the region of the observed variants. Also, customized visualization is enabled in the linked page “visualization options”, e.g. for selected chromosomal regions and grouping by subsets or studies. The lower panel is organized in four sections. 1) the “Result” tab (Figure 3F) shows the genome-wide CNA by the percentage of samples with yellow (+) as CN gain and blue (-) as CN loss. Below the CNA plot is a table showing the list of subsets as defined by ICD-O-3 and NCIt Ontology terms sorted by frequency of matched samples within that subset. 2) the “Biosamples” tab (Figure 3G) shows information of matched biosamples, i.e. description, classifications and external identifiers. The table can be downloaded as json or csv format. The further detail of the biosample can be accessed by clicking the biosample id. 3) The “Biosamples Map” tab (Figure 3H) shows a world map with the matched geological locations highlighted. 4) the “Variants” tab (Figure 3I) shows the variant “digest” (concatenated format with chromosome, start, end position and type of the CNA) and its corresponding biosample and callset. Likewise, the table can be downloaded as json or csv format.

**Fig 3.**
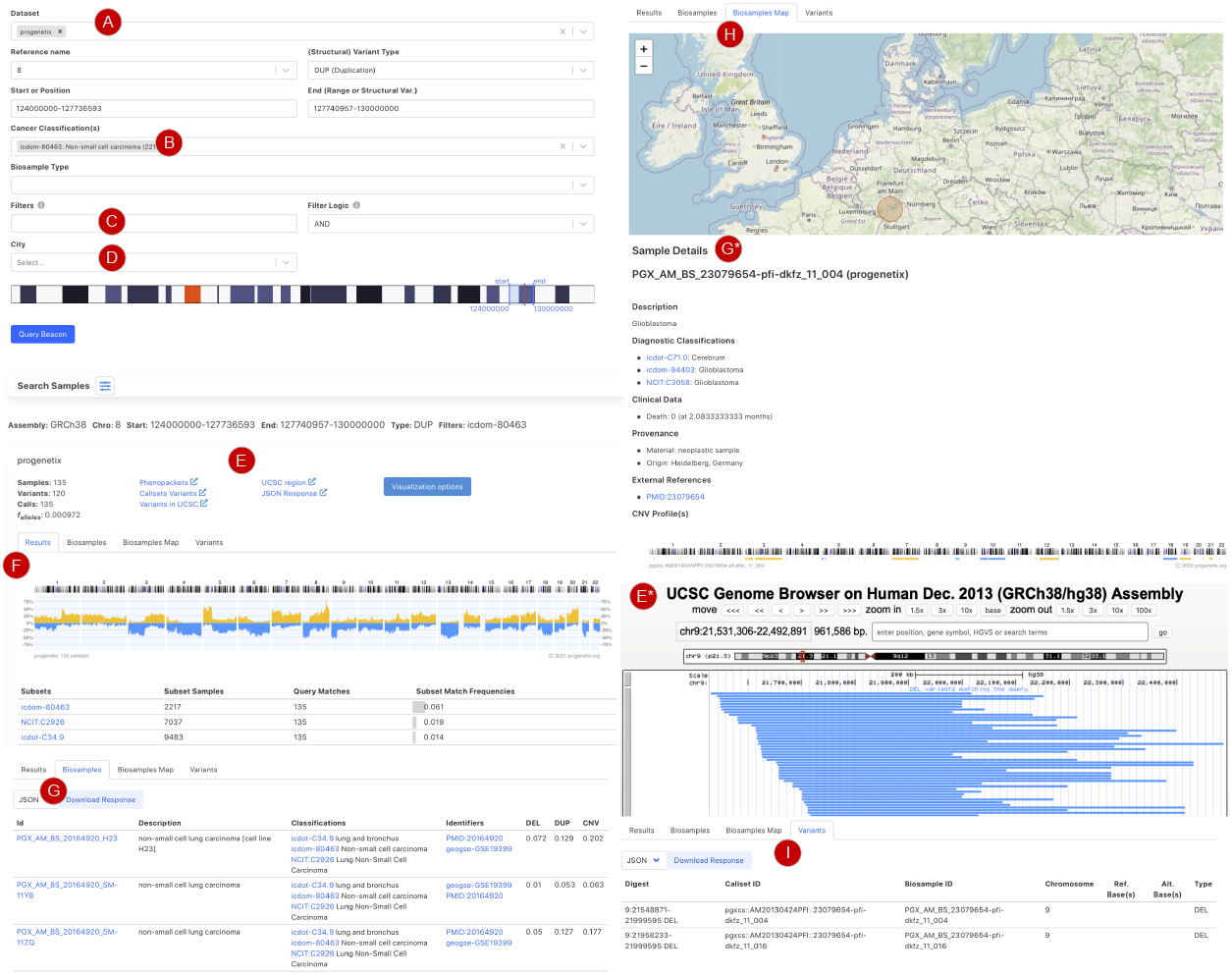
Beacon-style query using fuzzy ranges to identify biosamples with variants matching the CNA range. This example queries for a continuous, focal duplicaton covering the complete MYC gene’s coding region with <= 6Mbp in size. A: Filter for dataset. B: filter for cancer classification (NCIt and ICD-O-3 ontology terms available), C: additional filter, e.g. cellosaurus D: additional filter for geographic location. E: external link to UCSC browser to view the alignment of matched variants; F: cancer type classification sorted by frequency of the matched biosamples present in the subset; G: list of matched biosamples with description, statistics and reference. More detailed biosample information can be viewed through *id* link to the sample detail page. H: matched variants with reference to biosamples can be downloaded as json or csv format.

Figure 4 shows the additional functional interfaces and services provided by the Progenetix project. Users can search for publications or studies by publication title, author names or the geographic location of the research center. Then, navigation extends to the summary of publications with the number of samples catalogued by technology and availability in database as well as options to visualize the associated samples (Figure 4A). Users can also access samples from the NCIt hierarchical tree or other classification systems (e.g. ICD-O, UBERON) to select a subset of cancer types for summary statistics and visualization (Figure 4B). Alternatively, users can also upload their own data for single or multiple samples to visualize genome-wide CNA (Figure 4C).

**Fig 4.**
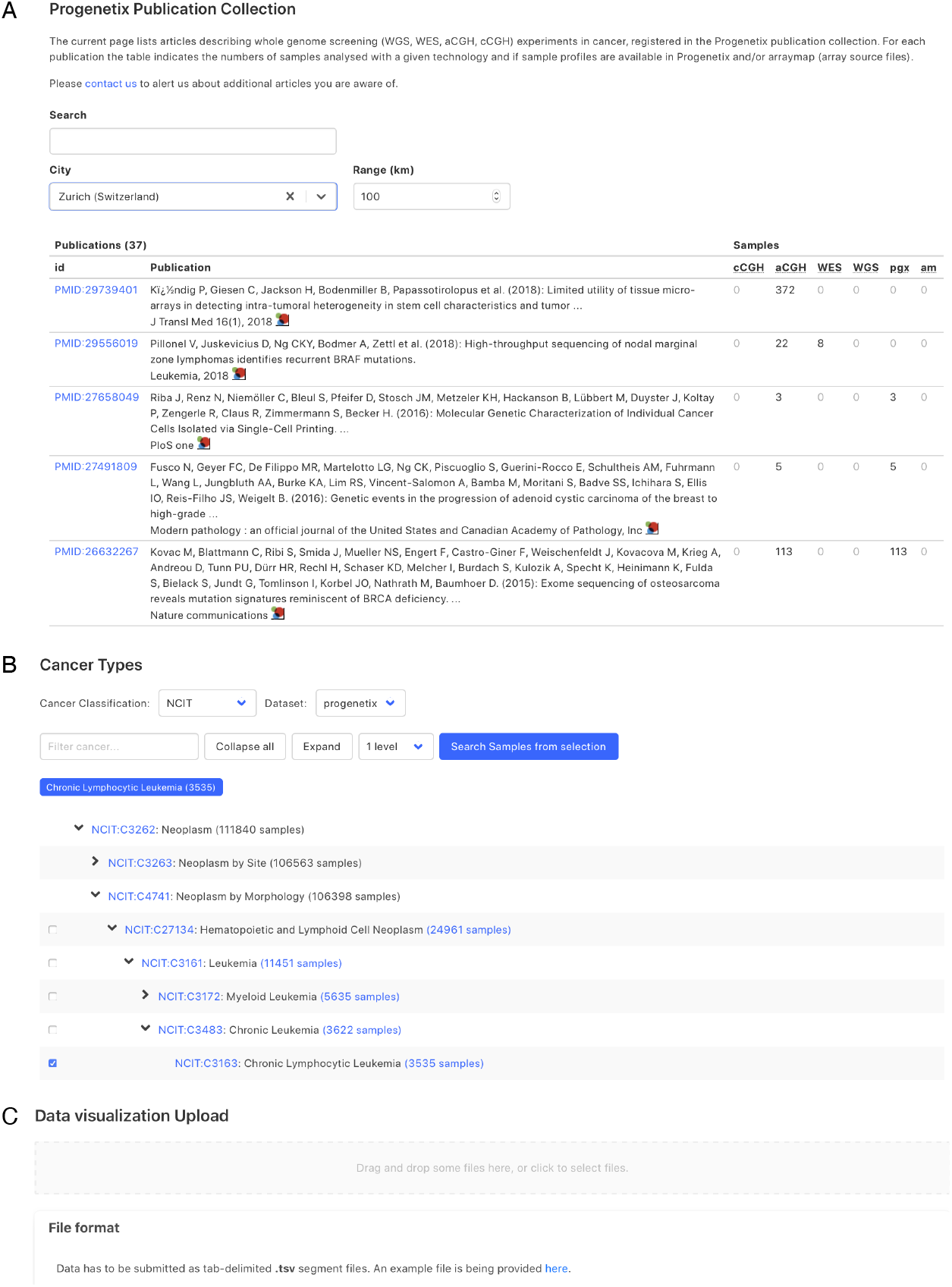
Demonstration of further functionality pages: A. Publication search; B. NCIT hierarchical tree navigation. A: Cancer genomics-associated publications are recorded with number of samples stratified by technology used. The publications can be filtered by keywords; B: Part of the sample subsets contained in Progenetix under the hierarchical NCIt classification tree. It allows for selection of sample subsets at different levels; C: User can upload custom segment files for data visualization.

## Other improvements

### Genome version update

All samples have been updated to Genome Reference Consortium Human Build 38 (GRCh38). The process has been completed in a step-wise manner. Preferably, for samples with available probe-specific array data, either GRCh38 mapped platform data files were used for re-processing of the original files; alternatively, a lift-over of the probe data and subsequent re-segmentation was performed. For those cases where only called CNA data had been collected we applied our recently published “segment-liftover” tool [19] for the efficient re-mapping of continuous segments. Overall, more than 99.99% of probes and more than 99% of segments could be recovered successfully.

### Cell line collection

Cancer cell lines are important models for understanding the molecular mechanisms of malignant diseases and have a prominent role in pharmacological screening procedures. Besides the primary tumor data, the Progenetix data collection also includes genomic profiling experiments using *in vitro* models. Recently, we introduced a systematic update of cell line annotations based on *Cellosaurus*, a comprehensive knowledge resource on cell line data with extensive annotations and mappings to a variety of classifications and ontologies [50]. We meticulously assigned Cellosaurus IDs for the cancer cell line samples as well as the ICD-O morphology and topography codes based on the NCIt term annotated by Cellosaurus. At this time, Progenetix includes a total of 5764 samples corresponding to 2162 different cancer cell lines, representing 259 different cancer types (NCIt). While so far we provide the option to search for cell lines by applying a “cellosaurus” filter either in the web interface (e.g. “cellosaurus:CVCL_0030” for *HeLa* cell line samples) or in the API query, work on a dedicated cell line data access tool is under way.

## Conclusion

The Progenetix resource provides an extensive collection of oncogenomic data with a focus on individual genome-wide CNA profiles and the use of modern ontologies and data schemas to render curated biological and technical metadata, as well as thorough references to external repositories and annotation resources. Through aggregation of data from thousands of individual research studies as well as several consortium derived collections, to our knowledge Progenetix database currently constitutes the largest public, freely accessible resource for pre-computed CNA profiles and associated phenotypic information and additional metadata dedicated to cancer studies. While the application of uniform genomic data formats and a benchmarked data processing pipeline minimizes biases from separate studies, the forward-looking implementation of emerging ontology standards facilitates the integrative and comparative analysis across a vast range of cancer types. The tight integration with GA4GH product development and standardization processes guarantees the compatibility with emerging data federation approaches and widest re-utilization of the resource’s data. For the future, besides the continuous maintainance and expansion of the existing data types, we will work towards enhancing clinical and diagnostic annotation, expanding cross-database references and the types of genomic variant data as well as active data sharing and integration through networked services and platforms.

## Supplementary Information

**Supplementary Figure 1**: The genome-wide CNV landscape of samples in the 51 NCIt categories. For each category, we randomly sampled around 150 genome profiling data and plotted the percentage of CN gain or loss across the genome. Yellow (+) indicates copy gain and Blue (-) indicates copy loss. The genomic profiles were ordered with hierarchical clustering with Euclidean distance and median linkage. NOS: not otherwise specified.

**Supplementary Figure 2**: The genome-wide LOH landscape of samples in the 51 NCIt categories. For each category, we randomly sampled around 100 genome-wide LOH profiles and plotted the percentage of samples harboring LOH in regions across the genome in the (-) direction, as an indication for uniparental allelic loss. The genomic profiles were ordered with hierarchical clustering with Euclidean distance and median linkage. NOS: not otherwise specified.

**Supplementary Table 1**: The number of samples belonging to the 51 NCIt summary terms in Progenetix.

**Supplementary Table 2**: The mapping table between the 1000 Genomes reference population labels and HANCESTRO ontology terms.

## Acknowledgments

Work on the Progenetix Beacon implementation was supported by ELIXIR and through the BioMedIT Network project of SIB Swiss Institute of Bioinformatics in the realm of the Swiss Personalized Health Network. We’d like to thank Amos Bairoch for support with the cell line annotations. Improvements in data annotation concepts were highly influenced through the GA4GH community.

