## Supplementary figures and images for "The Progenetix oncogenomic resource in 2021"

### Supplementary Figure 1

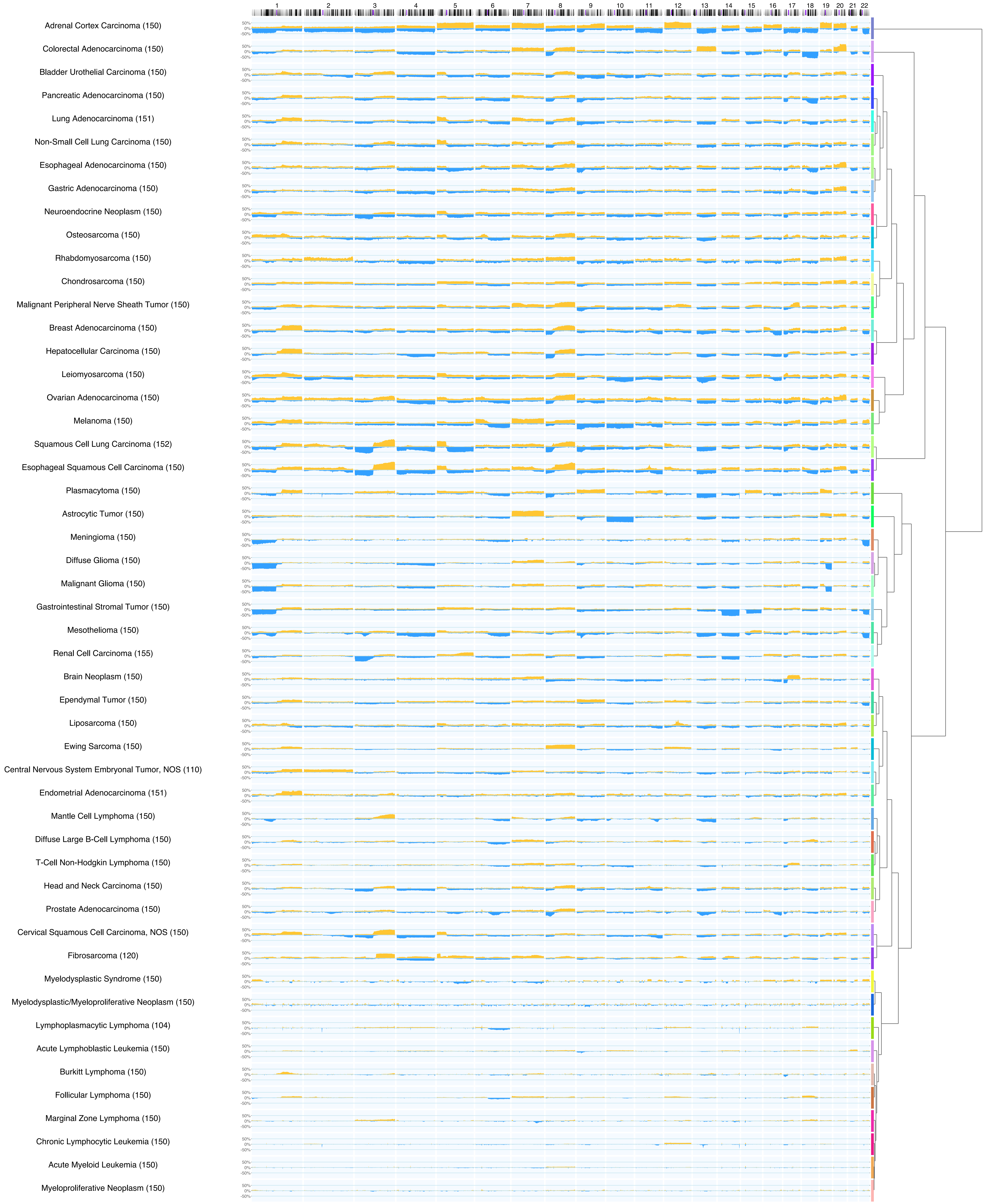

### Supplementary Figure 2

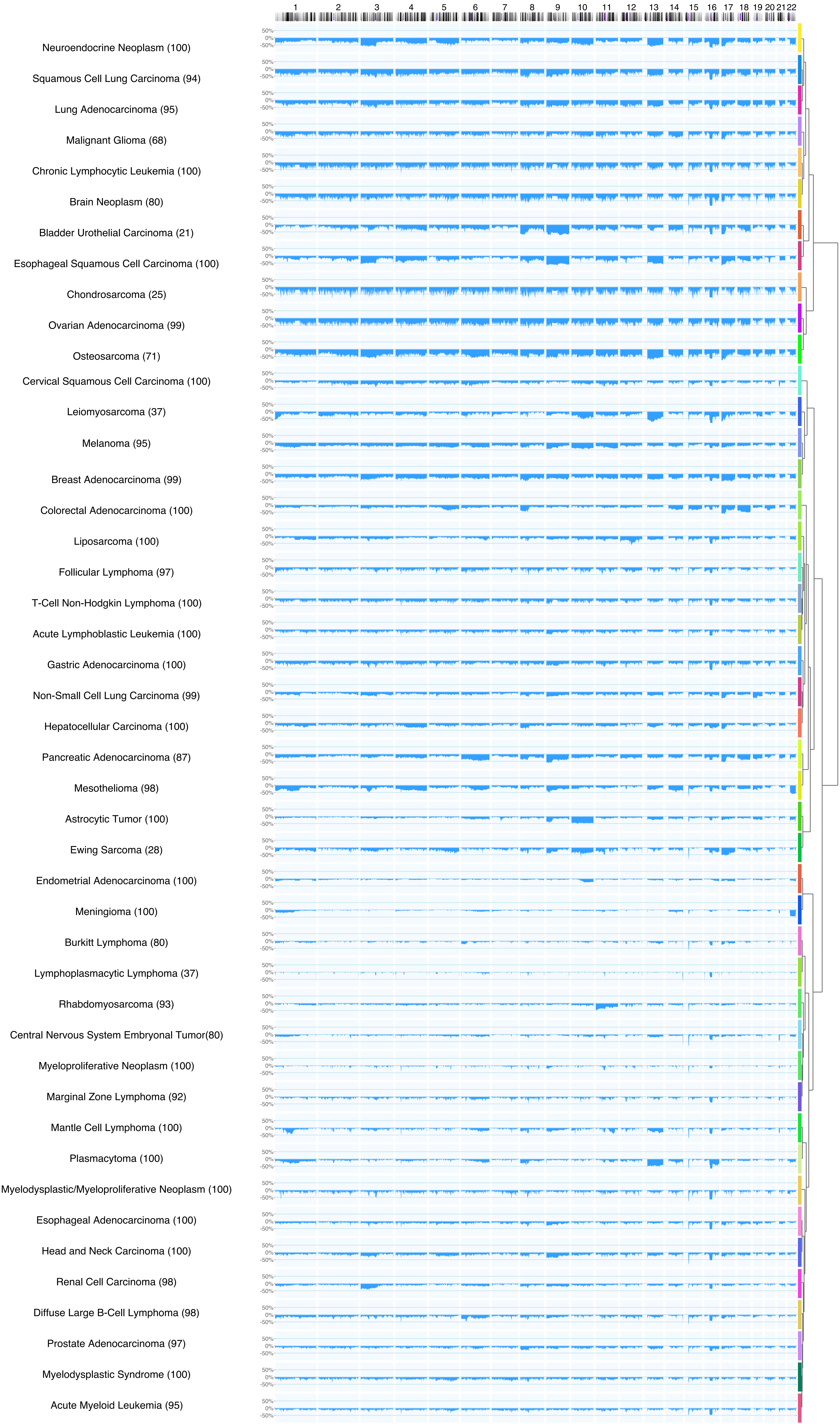
